# The Structural Flexibility of MAD1 Facilitates the Assembly of the Mitotic Checkpoint Complex

**DOI:** 10.1101/2022.06.29.498198

**Authors:** Chu Chen, Valentina Piano, Amal Alex, Simon J. Y. Han, Pim J Huis In ’t Veld, Babhrubahan Roy, Andrea Musacchio, Ajit P. Joglekar

## Abstract

The spindle assembly checkpoint (SAC) safeguards the genome during cell division by generating an effector molecule known as the Mitotic Checkpoint Complex (MCC). The MCC comprises two subcomplexes, and during its assembly, formation of the CDC20:MAD2 subcomplex is the rate-limiting step. Recent studies show that the rate of CDC20:MAD2 formation is significantly accelerated by the cooperative binding of CDC20 to SAC proteins MAD1 and BUB1. However, the molecular basis for this acceleration is not fully understood. Here, we demonstrate that the structural flexibility of MAD1 at a conserved hinge near the C-terminus is essential for catalytic MCC assembly. This MAD1 hinge enables the MAD1:MAD2 complex to assume a folded conformation *in vivo*. Importantly, truncating the hinge reduces the rate of MCC assembly *in vitro* and SAC signaling *in vivo*. Conversely, mutations that preserve hinge flexibility retain SAC signaling, indicating that the structural flexibility of the hinge, rather than a specific amino acid sequence, is important for SAC signaling. We summarize these observations in a “knitting” model that explains how the folded conformation of MAD1:MAD2 promotes CDC20:MAD2 assembly.

---

During mitosis, a parent cell divides into two genetically identical daughter cells. To achieve this, the duplicated chromosomes in the parent cell must be equally distributed into the two daughter cells. The spindle assembly checkpoint (SAC) serves as a surveillance mechanism to ensure that duplicated chromosomes are stably attached to spindle microtubules through an adaptor structure named the kinetochore. Kinetochores lacking end-on microtubule attachment activate the SAC to prevent premature anaphase onset and avoid chromosome missegregation. The effector molecule generated upon SAC activation is the Mitotic Checkpoint Complex (MCC). The MCC consists of two subcomplexes: BUBR1:BUB3 and CDC20:MAD2 ^1, 2^. It inhibits the E3 ubiquitin ligase anaphase-promoting complex/cyclosome (APC/C) ^3-5^. APC/C ubiquitinates Cyclin B1, a key mitosis regulator, thereby targeting it for proteasome-mediated degradation ^6-8^. Inhibition of the APC/C suppresses the degradation of Cyclin B1, which in turn delays anaphase onset.

The formation of the CDC20:MAD2 dimer has been identified as the rate-limiting step in the assembly of the MCC ^9, 10^. This biochemical step is catalyzed by other checkpoint proteins, including the MAD1:MAD2 complex and the BUB1:BUB3 complex, that recruit the MCC subunits and facilitate their interaction. A crucial aspect of the reaction that allows MAD2 to bind CDC20 is the conversion of MAD2 from an open conformation (O-MAD2) to a closed conformation (C-MAD2) ^11-14^. During this conversion, the C-terminal “safety belt” of MAD2 embraces the flexible MAD2-interacting motif (MIM) of CDC20 ^2, 13^. Purified monomeric O-MAD2 spontaneously converts into C-MAD2 at 30 °C *in vitro* with kinetics that are orders of magnitude slower than expected to support robust CDC20:MAD2 formation during mitosis ^15^. In a reconstituted reaction *in vitro*, MAD1:MAD2 and BUB1:BUB3 were shown to dramatically accelerate the assembly of the CDC20:MAD2 complex, suggesting that they are the catalysts in the assembly reaction ^10, 16^. The “MAD2 template model” ^14^ argues that the conformational switch is facilitated by the dimerization between one C-MAD2 (bound to MAD1’s MIM in the MAD1:MAD2 complex) and a cytosolic O-MAD2 that undergoes the conformational switch to bind CDC20. Furthermore, two recent studies show that the docking of CDC20 on multiple interfaces on MAD1 and BUB1 enables spatio-temporal coupling of the MAD2 conformational switch with its binding to CDC20 thereby overcoming the rate-limiting step and accelerating MCC assembly ^16, 17^. The exact molecular mechanism of this coupling, however, remains to be elucidated.

In this paper, supported by modeling of the MAD1:MAD2 complex, we hypothesize that efficient CDC20:MAD2 formation may require a folded conformation of the “MAD1 C-terminal region”. We define the MAD1 C-terminal region as spanning residues 485-718 including the Mad1 C-terminal domain (Mad1-CTD) known to be essential for catalysis ^10, 13, 17-21^. In agreement with this hypothesis, fluorescence-lifetime imaging (FLIM) suggests that the C-terminal hinge of MAD1 enables the MAD1:MAD2 complex to take a folded conformation *in vivo*. Importantly, disrupting the structural flexibility of MAD1 by removing the hinge impairs the rate of MCC assembly *in vitro* and the SAC signaling activity *in vivo*. Mutating this region while keeping its flexibility maintains the SAC signaling activity, indicating that the structural flexibility (rather than the primary sequence specificity) of MAD1 is important to the SAC. We propose a “knitting model” that describes how the MAD2 conformational switch is coupled to the formation of CDC20:MAD2, which is key for rapid activation of the SAC in living cells.

## The MAD1:MAD2 complex may assume a folded conformation *in vivo*

The MAD1:MAD2 complex is a 2 : 2 heterotetramer. Prior studies have defined the structures of two non-overlapping, dimeric segments of the of the C-terminal region of this heterotetramer: one spanning residues 485-584 and complexed with two MAD2 molecules, and the other, termed as the Mad1-CTD, spanning residues 597-718 ^13, 18^. The SAC kinase MPS1 phosphorylates T716 within the RING finger-containing proteins, WD repeat-containing proteins, and DEAD-like helicases (RWD) domain at the C-terminus of MAD1. Upon phosphorylation, MAD1-CTD binds the BOX1 motif in the N-terminal region of CDC20 ^16, 21^, and this interaction is critical for MCC assembly ^10, 21, 22^. It likely facilitates the coupling of the MAD2 conformational switch with CDC20 binding. However, if we model the disordered N-terminus of human CDC20 as a simple 3-D random walk, the estimated root-mean-square distance from BOX1 (27–34) to MIM (129–133) is less than 4 nm. The worm-like chain model with a persistence length of 0.3–0.7 nm estimates the root-mean-square distance to be 4.5–7.4 nm ^23, 24^. On the other hand, the combined axial length from the MAD1 MIM to the RWD domain is over 12 nm, according to crystal structures of the two MAD1:MAD2 segments (Figure 1A, left panel) ^13, 18^. Therefore, the flexibility of the CDC20 N-terminus may not be sufficient to position the MIM of CDC20 proximally with respect to MAD2, and additional mechanisms may facilitate the efficient capture of the CDC20 MIM by MAD2.

**Figure 1.**
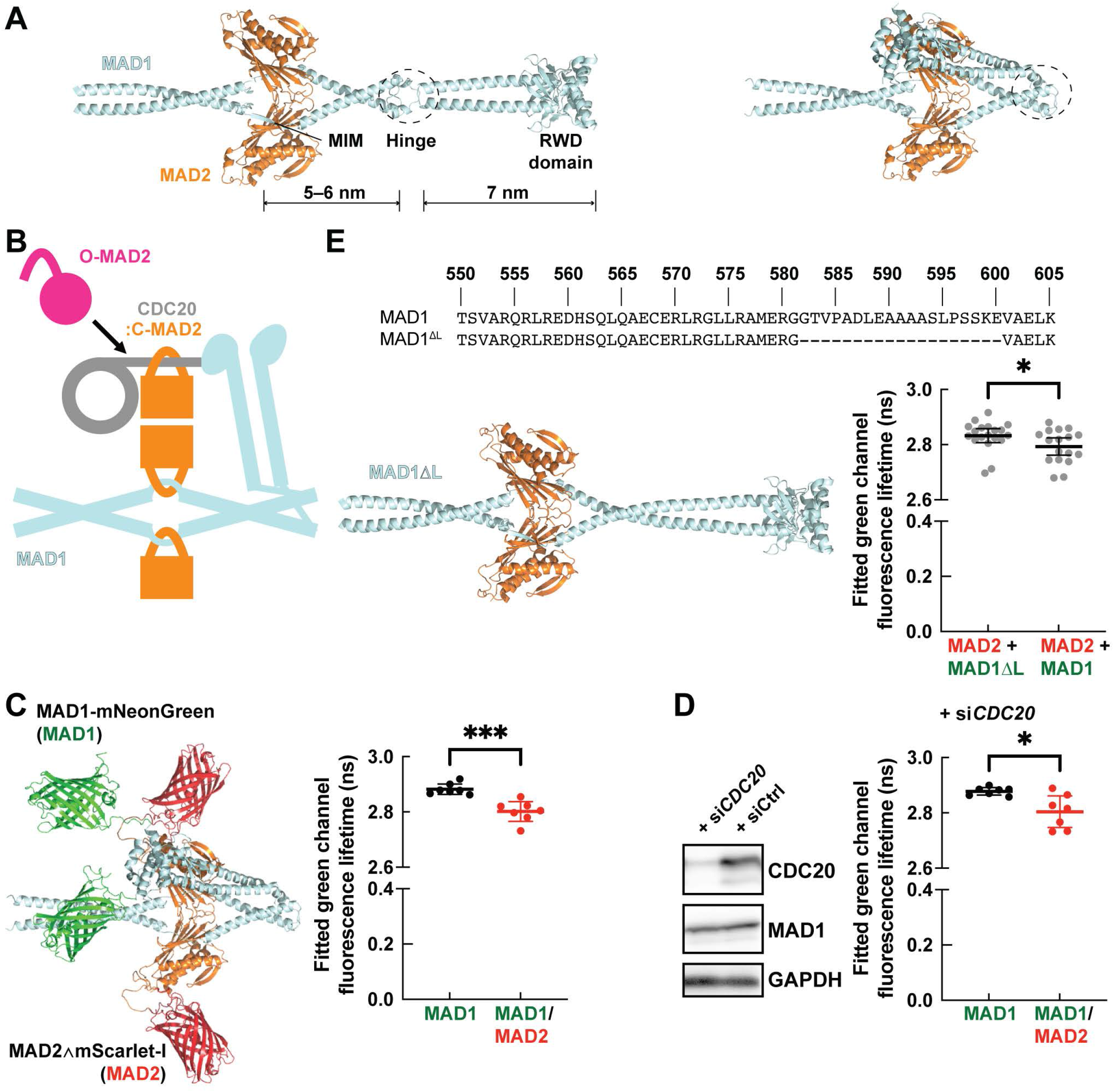
The MAD1:MAD2 complex can assume a folded conformation *in vivo* enabled by the hinge of MAD1. (A) Representative models of the core region of the MAD1:MAD2 complex were predicted by the ColabFold advanced algorithm. The complex can assume either an extended (left) or a folded (right) conformation. The hinge is circled out. These predicted structures agree with published crystal structures, from which labeled length measurements were taken (PDB IDs: 1GO4, ref. ^13^ and 4DZO, ref. ^18^). (B) A cartoon demonstrating how the folded conformation helps present the MIM of CDC20 to MAD2 that undergoes the conformational switch. The N-terminal region (containing BOX1 and the MIM) and C-terminal region (the WD40 fold) of CDC20 are represented by a light gray line and a light gray circle, respectively. The contact between CDC20’s N-terminal region and MAD1 represents direct physical interaction ^16, 21, 40^. All cartoons in this paper are not to scale. (C) The core region of the MAD1-mNG:MAD2∧mScarlet-I complex in a folded conformation predicted by the ColabFold advanced algorithm (left) and the average lifetime of MAD1-mNG in the *MAD1*-mNG genome-edited HeLa-A12 cell line or the *MAD1*-mNG/*MAD2*∧mScarlet-I genome-edited HeLa-A12 cell line (right). The total number of cells in each group *N* = 7. Results are representative of two independent experiments. (D) (Left) Unsynchronized HeLa-A12 cells were treated with si*CDC20* or a control siRNA for 2 d and probed for CDC20, MAD1, and GAPDH (loading control). (Right) Same as (C), except that cells were treated with si*CDC20*. (E) (Top) Partial sequence of human wild-type MAD1 or MAD1ΔL. (Bottom left) A representative model of the core region of the MAD1ΔL:MAD2 complex predicted by the ColabFold advanced algorithm. (Bottom right) The average lifetime of exogenous MAD1ΔL-mNG and MAD1-mNG in the *MAD2*∧mScarlet-I genome-edited HeLa-A12 cell line. The total number of cells in each group *N* ≥ 17. Results are pooled from four independent experiments. In (C) to (E), each dot represents a single cell. Mean values ±95% confidence intervals are overlaid. Unpaired *t*-tests with Welch’s correction are performed in Prism (GraphPad Software). The following symbols for p-values are used in this paper: ns (not significant, *p* ≥ 0.05), ∗(0.01 ≤ *p* < 0.05), ∗∗(0.001 ≤ *p* < 0.01), ∗∗∗(0.0001 ≤ *p* < 0.001), and ∗∗∗∗(*p* < 0.0001).

To gather possible clues, we used AlphaFold 2 ^25, 26^ to predict how the structurally known segments of MAD1 may be arranged. In addition to an extended conformation, this analysis predicted the existence of a folded conformation of MAD1 enabled by a flexible hinge spanning residues 582 and 600 (Figure 1A, right panel). We reasoned that the folded MAD1 conformation would permit the phosphorylated C-terminal RWD domains of MAD1 to approach the reaction center of the MAD1:MAD2 template complex where O-MAD2 is expected to undergo the conformational switch and bind CDC20 (Figure 1B). Interestingly, the primary sequence of the hinge region is not conserved from yeast to human (Figure S2B), but an interruption of the coiled-coil around this region appears to be universal (Figure S2C). According to AlphaFold 2 predictions, the flexibility of the hinge region enables MAD1 to assume a spectrum of conformations, from fully extended to folded (Figure 1A) ^25, 26^.

To test whether the MAD1:MAD2 complex assumes a folded conformation *in vivo*, we resorted to distance-sensitive Förster resonance energy transfer (FRET) assays. The folded conformation is expected to drastically reduce the distance between the RWD domain and the MIM of MAD1, and a correctly designed FRET sensor may be able to differentiate the folded conformation from the extended conformation (Figure 1A). Using CRISPR-Cas9-mediated genome editing, we fused the donor fluorophore mNeonGreen to the C-terminal end of MAD1 and inserted the acceptor fluorophore mScarlet-I ^27^ in the β5-αC loop of MAD2 (the exogenous protein is henceforth referred to as “MAD2∧mScarlet-I”; Figures S1A and S1C) ^28^. This strategy positions the acceptor fluorophore away from known functional interfaces of MAD2 (including the MAD2 homodimerization interface, the safety belt, and the interface between MAD2 and BUBR1 in the MCC) ^2, 14, 28, 29^. This strategy also takes into account our unpublished observations that in the budding yeast *Saccharomyces cerevisiae*, neither N-nor C-terminally tagged Mad2 supports SAC signaling. In the extended conformation, the distance between the donor and acceptor will be larger than 10 nm, allowing minimal FRET between the donor and the acceptor ^30^. Conversely, in the folded conformation, the distance between the donor and the acceptor will be reduced, increasing the efficiency of FRET between the donor and the acceptor (Figure 1C, left panel).

We first tested that in the budding yeast, exogenous internally-tagged Mad2 supports the SAC activity in the *mad2Δ* background (Figure S1B). Next, we confirmed the expression of full-length MAD2∧mScarlet-I in the heterozygous *MAD2*∧mScarlet-I genome-edited HeLa-A12 cell line, wherein the expression level of either BUBR1 or CDC20 was not affected (Figure S1D). Internally tagged MAD2 partially restores the SAC signaling activity in HeLa-A12 cells when endogenous MAD2 is knocked down via RNA interference (Figure S1E).

Using FLIM ^31^, we quantified a FRET efficiency of about 3% between MAD1-mNG and MAD2∧mScarlet-I at the interphase/prophase nuclear pore complex (NPC), where MAD1:MAD2 resides during interphase, in the heterozygous *MAD1*-mNG, *MAD2*∧mScarlet-I genome-edited HeLa-A12 cell line (Figure 1C, right panel). We measured FRET at the interphase/prophase NPC to facilitate data collection by the line-scanning confocal microscope and to reduce the interference of potential intermolecular FRET between a donor from one MAD1:MAD2 complex and an acceptor from another nearby complex, which is expected at the corona of a signaling kinetochore. This FRET persists even when *CDC20* is knocked down by RNAi (Figure 1D), suggesting that it is intrinsic to the MAD1:MAD2 complex. It should be noted that two experimental details contribute to the low FRET efficiency observed. First, only half of MAD1 and MAD2 are fluorescently labeled. Second, the combined size of the fluorescent proteins and the flexible linkers used to fuse them to MAD1 and MAD2 adds a significant distance to the actual separation between the two proteins ^30^.

To reinforce these observations, we designed a MAD1 mutant (henceforth referred to as MAD1ΔL) wherein the hinge (582–600) was deleted. This deletion preserves the heptad repeat periodicity of the upstream and downstream coiled-coils predicted by MARCOIL and DeepCoil2 (data not shown) ^32-34^. For the resulting hinge-deleted MAD1 mutant, AlphaFold 2 predicted an uninterrupted and fully extended coiled-coil (Figure 1E, bottom left panel). The FRET efficiency of the mutant was reduced by half (to 1.5%). Even though there is some residual FRET between MAD1ΔL-mNG and MAD2∧mScarlet-I (Figures 1C and 1E), these results support that the structural flexibility of the C-terminus of MAD1 enabled by the hinge facilitates folding of the MAD1:MAD2 complex *in vivo*.

## MAD1’s hinge is important to the rate of MCC assembly *in vitro*

To test the role of the structural flexibility of MAD1 in the assembly of the MCC, we purified recombinant MAD1:MAD2 and MAD1ΔL:MAD2 and compared their functionality in the previously established MCC FRET-sensor-based assays ^10, 16^. Importantly, the complexes appeared stable and properly folded (Figure S2A).

Deletion of MAD1’s hinge causes a moderate but reproducible decrease in the rate of MCC assembly compared to the wild-type (Figure 2B), indicating that the hinge is important to maximize the rate of MCC assembly *in vitro*. The rate difference between MAD1:MAD2 and MAD1ΔL:MAD2 relied on the presence of BUB1:BUB3 (Figure 2C). More specifically, the rate difference required a functionally intact BUB1:BUB3 complex to interact with MAD1:MAD2, because the BUB1ΔCM1 mutant that prevents this interaction erased the difference (Figure S2D).

**Figure 2.**
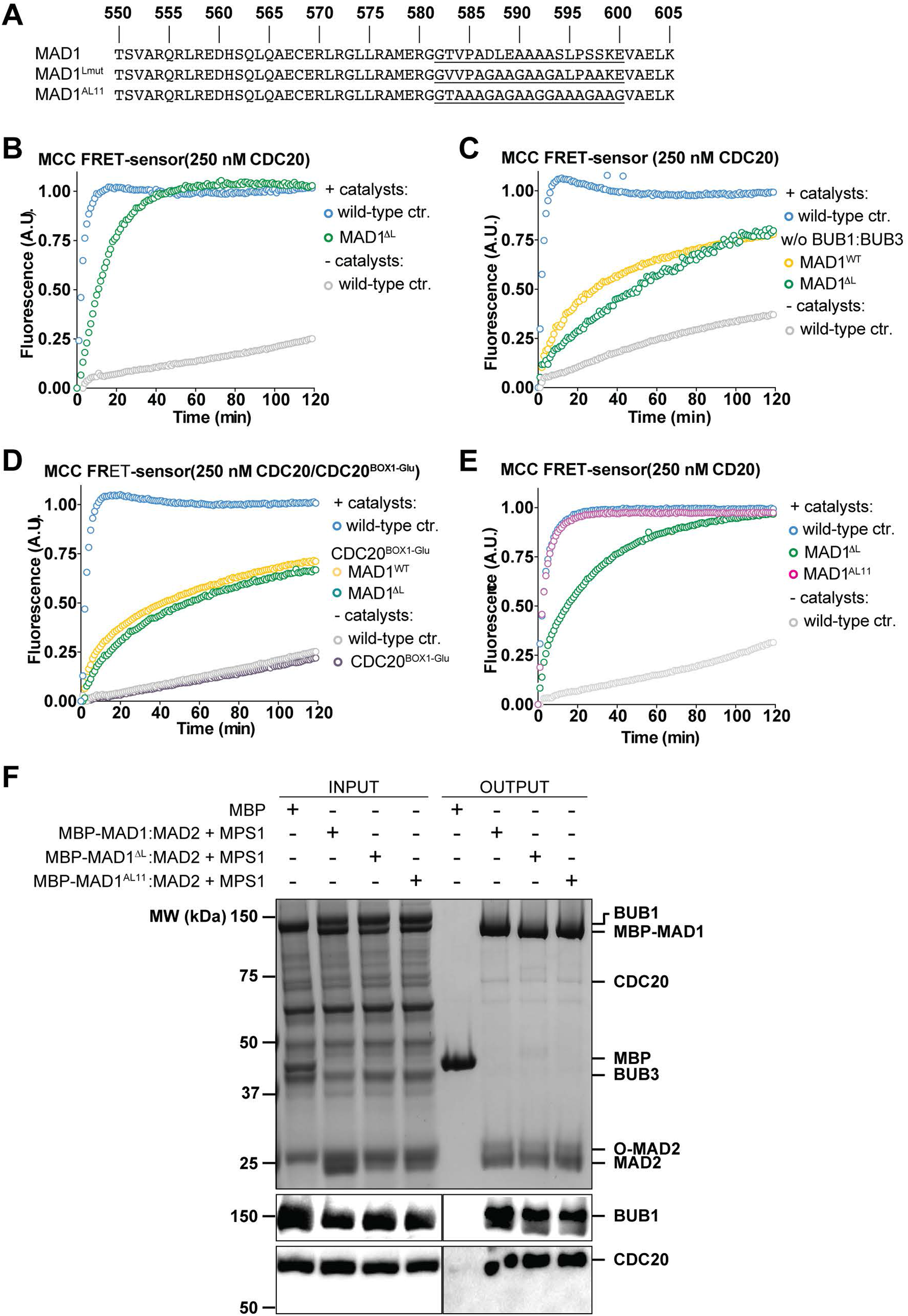
The rate of MCC assembly is lower in the presence of MAD1ΔL than in the presence of wild-type MAD1 *in vitro*. (A) Partial sequence of wild-type or mutant human MAD1. The hinge (582–600) is underlined. (B) The addition of MBP-MAD1ΔL:MAD2 (green) causes a moderate decrease in the rate of MCC assembly compared to the wild-type (blue). (C–D) MBP-MAD1:MAD2 (yellow) and MBP-MAD1ΔL:MAD2 (green) have similar MCC assembly rates (C) in the absence of BUB1:BUB3 or (D) when CDC20^BOX1-Glu^ is used in the reaction instead of wild-type CDC20. (E) MBP-MAD1(AL11):MAD2 (magenta) can promote MCC assembly *in vitro* similarly to wild-type MBP-MAD1:MAD2 (blue). In (B) to (E), curves report single measurements are representative of at least three independent technical replicates. The *y*-axis represents the normalized emission intensity of the acceptor. Prism was used for data analysis and visualization. (F) MBP or MBP-MAD1(wild-type or mutant):MAD2 is immobilized on amylose beads and serves as baits to pull down preys including O-MAD2 (a V193N mutant that stabilizes MAD2 in the open conformation ^28^), MPS1-phosphorylated BUB1:BUB3, and CDC20. From top to bottom: a Coomassie-stained SDS-PAGE gel, an immunoblot detecting BUB1, and an immunoblot detecting CDC20.

Manifestation of a rate difference between MAD1:MAD2 and MAD1ΔL:MAD2 also relied on the interaction of CDC20 with MAD1, as it was abolished by mutation of the BOX1 motif of CDC20 (Figure 2D). Collectively, these observations suggest that flexibility enabled by the hinge region allows MAD1:MAD2 to interact more productively with BUB1 and CDC20 during the catalytic conversion that promotes MCC assembly. In a solid phase binding assay with immobilized MAD1:MAD2, we found that the binding of O-MAD2, BUB1, and CDC20 was not overtly affected by the hinge-deletion mutation (Figure 2F). We conclude that the role of the structural flexibility of MAD1 in the rate of MCC assembly *in vitro* is critical to the appropriate spatial association of BUB1, CDC20, and MAD1:MAD2 ^10, 16, 17^ ; see Discussion).

## The C-terminal hinge of MAD1 is important to the SAC signaling activity *in vivo*

Next, we sought to determine whether the C-terminal hinge of MAD1 is important for SAC signaling *in vivo*. We integrated the expression cassette of either MAD1-mNG or MAD1ΔL-mNG into the genome of HeLa-A12 cells using Cre-*lox* recombination-mediated cassette exchange (RMCE) ^35-37^. We then knocked down endogenous MAD1 in these cells using two siRNAs that target the 3’-UTR of *MAD1* ^38^ (henceforth collectively referred to as si*MAD1*’s) and induced the expression of MAD1(WT/ΔL)-mNG (si*MAD1*-resistant due to the lack of the endogenous 3’-UTR) by doxycycline. Our genome-edited *MAD1*-mNG HeLa-A12 cell line served as the reference for the endogenous level of MAD1 in live-cell fluorescence imaging. When quantifying the phenotypes of the knock-in/knock-down treatments, we ensured that the kinetochore recruitment of the MAD1 mutants used was comparable to the recruitment of *MAD1*-mNG in the genome-edited HeLa-A12 cells (see Methods).

Cells with less than 10% of the physiological level of MAD1 generally retained a robust checkpoint response in 100 nM nocodazole that could not be weakened by increasing the dosage of si*MAD1*’s (Figure S3A and S3B). Nonetheless, SAC signaling activity was crippled, as the depletion caused MAD1-depleted cells to leave mitosis at least two hours earlier than the untreated control (Figure 3A). In this context, however, expression of MAD1ΔL-mNG resulted in a dominant-negative effect that considerably shortened the mitotic arrest. For comparison, wild-type MAD1-mNG restored the SAC signaling activity to levels observed in the negative control (Figure 3A). We reasoned that the dominant-negative effects of MAD1ΔL-mNG reflect its dimerization with the residual endogenous MAD1 and consequent restriction of its structural flexibility. Indeed, an AlphaFold 2 structural prediction of MAD1:MAD1ΔL suggested that the hinge region of wild-type MAD1 cannot adopt the folded conformation when facing the stiff continuous α-helix of the MAD1ΔL counterpart (Figure 3B). To test this experimentally, we pulled down doxycycline-induced MAD1(wild-type/ΔL)-mNG from lysates of HeLa-A12 cells in which endogenous MAD1 was not knocked down. We found that endogenous MAD1 was pulled down both by MAD1-mNG and by MAD1ΔL-mNG, but not by mNeonGreen alone (Figure S3D). We further confirmed that MAD1ΔL-mNG did not cause defects in the localization of the MAD1ΔL:MAD2 complex (Figure S3C) or the expression of BUBR1, CDC20, or BUB3 (Figure S3B). Therefore, although the results of our knockdown-rescue experiments were hindered by the incomplete knockdown of the endogenous MAD1, all evidence combined suggested that the hinge of MAD1 is critical for the SAC.

**Figure 3.**
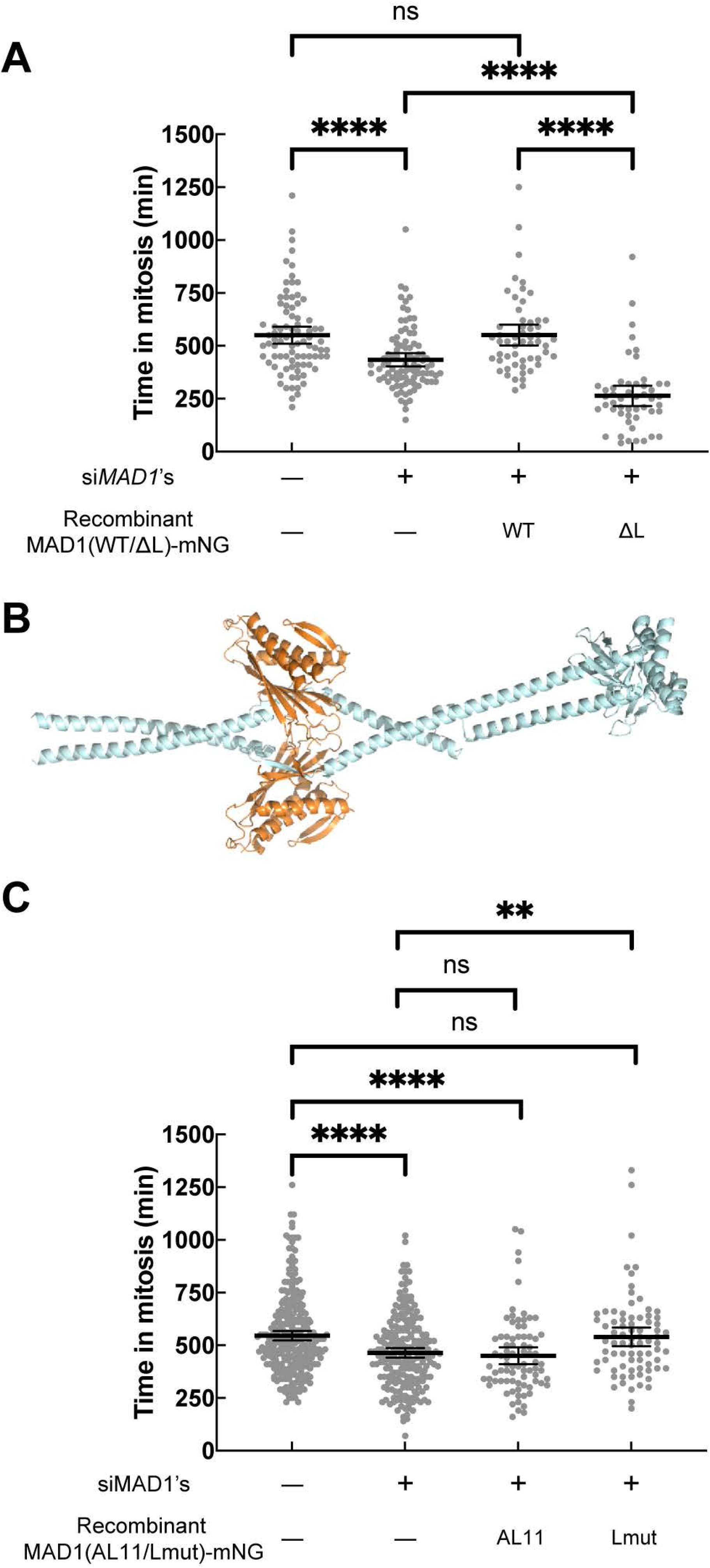
The structural flexibility provided by the hinge of MAD1 is critical to the SAC signaling activity *in vivo*. (A) The first two columns on the left used the *MAD1*-mNG genome-edited HeLa-A12 cell line which served as a reference for the endogenous level of MAD1 (see Methods). *In situ* tagging of MAD1 did not affect the 3’-UTR which si*MAD1*’s target. The effectiveness of si*MAD1*’s against the *MAD1*-mNG allele was confirmed by the greatly diminished green channel fluorescence signal (data not shown). The two columns on the right used HeLa-A12 cell lines treated with si*MAD1*’s and induced to express exogenous MAD1-mNG or MAD1ΔL-mNG. Each dot represents a cell (*N* ≥ 50 in each group). (B) In the predicted structure of the core region of the MAD1:MAD1ΔL heterodimer (in complex with MAD2, using the ColabFold advanced algorithm), the hinge of the wild-type copy introduces a bulge but the overall conformation is extended due to the stiffness of the now fused α-helix of MAD1ΔL. (C) As in (A), the first two columns on the left used the *MAD1*-mNG genome-edited HeLa-A12 cell line which served as a reference for the endogenous level of MAD1. The two columns on the right used HeLa-A12 cell lines treated with si*MAD1*’s and induced to express exogenous MAD1(AL11)-mNG or MAD1(Lmut)-mNG. Each dot represents a cell (*N* ≥ 75 in each group). In (A) and (C), results were pooled from at least two technical repeats. The mean value ± the 95% confidence interval of each group is overlaid. Unpaired *t*-tests with Welch’s correction are performed in Prism.

## MAD1(Lmut) can fully support the SAC signaling activity

The observation that the hinge encompassing residues 582–600 of MAD1 is important for SAC signaling *in vivo* may have alternative explanations. For instance, it is known that S598 can be phosphorylated by MPS1 *in vitro* ^21^, and we cannot exclude that the hinge of MAD1 is required for regulated, but unknown, protein-protein interactions important to the SAC. To distinguish among these possibilities, we reasoned that replacing the hinge with an equally flexible region of a diverged sequence should prevent sequence-specific physical interactions with putative binding partners while preserving MAD1’s ability to adopt the folded conformation. Therefore, we tested two different artificial flexible hinges, “AL11” and “Lmut”, as a replacement for the original hinge segment (Figure 2A). Both replacements consist of 19 amino acid residues as the original hinge. AL11 is a previously characterized flexible linker composed of eleven alanine residues, seven glycine residues, and one threonine residue ^39^. In Lmut, serine and threonine residues of the original segment are mutated into alanine and valine residues, respectively. The amino acids between the two prolines consist mostly of alanine-glycine-alanine repeats while the proline residues themselves and their N-terminal neighboring residues are preserved. Both MAD1(AL11) and MAD1(Lmut) are predicted to have a coiled-coil propensity profile similar to that of the endogenous MAD1 (Figure S4).

We observed that MAD1(AL11):MAD2 had the same MCC assembly activity as the wild-type complex in our *in vitro* assay (Figure 2E). We were unable to purify recombinant MAD1(Lmut):MAD2, possibly because of instability during protein purification introduced by the mutation. Both mutants were correctly expressed in HeLa cells (Figure S4B). Furthermore, in cells treated with si*MAD1* and expressing MAD1(AL11)-mNG, SAC signaling appeared slightly weaker than in cells expressing wild-type MAD1, while MAD1(Lmut)-mNG fully restored the SAC signaling activity (Figure 3C). We conclude that both MAD1 constructs with an artificial hinge are largely or completely checkpoint proficient, contrary to MAD1ΔL. These observations suggest that the primary function of the hinge is providing structural flexibility rather than mediating unspecified protein-protein interactions.

## Discussion

Here, we identified a previously unrecognized molecular mechanism that helps overcome the kinetic barrier associated with the binding of MAD2 and CDC20. A folded conformation of MAD1 positions the MIM of CDC20 and MAD2 closely, facilitating the assembly of the CDC20:MAD2 heterodimer. In a complementary study ^40^, Fischer and colleagues demonstrate that the CM1 of human BUB1 and the α1 helix of CDC20, which precedes BOX1, interact in a tripartite 1:1:2 complex with the RLK motif of MAD1. Thus, collectively, CDC20 establishes multiple interfaces with the catalysts BUB1 and MAD1:MAD2, and these interactions likely position the CDC20 MIM for its efficient capture by MAD2. Switching back to an extended conformation may break the avidity, thereby releasing assembled CDC20:MAD2 into the cytosol. We use the “knitting” analogy to describe this model (Figure 4), as the two MAD1 functional regions connected by the hinge switch their relative positioning and work coordinately like two knitting needles to “entangle” CDC20 and MAD2.

**Figure 4.**
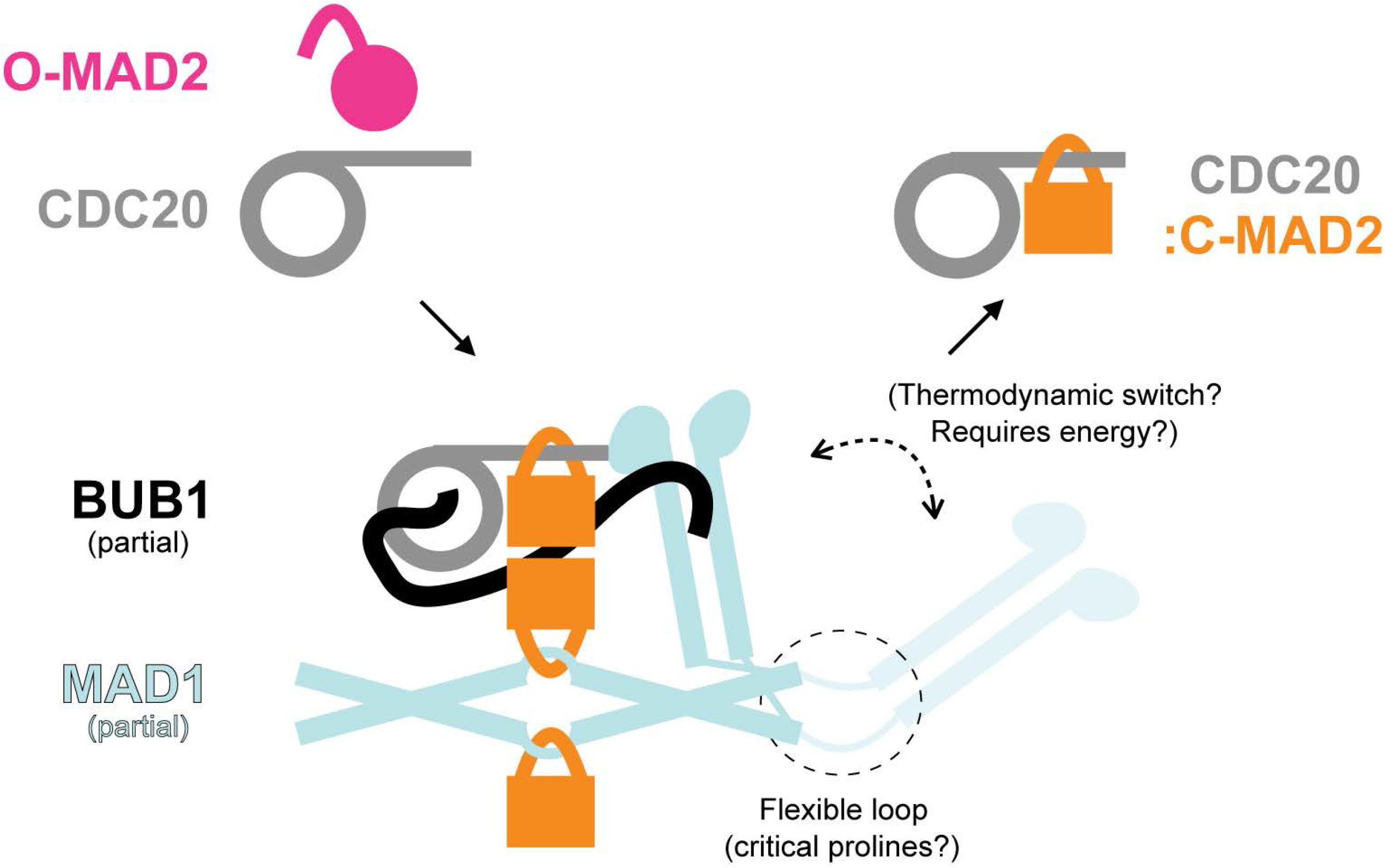
A cartoon of the “knitting” model. The structural flexibility of MAD1 facilitates the spatio-temporal coupling of the MAD2 conformational switch and the assembly of CDC20:MAD2. The two solid black arrows indicate the formation and release of CDC20:MAD2, respectively. According to Figures 2C and S2D, the difference in the MCC assembly rate (comparing MAD1 with MAD1ΔL) relies on the interaction between MAD1 and BUB1. Therefore, this cartoon of our model also incorporates BUB1 and highlights the following protein-protein interactions involving BUB1: (1) T461-phosphorylated BUB1 CM1 interacts with MAD1’s consensus RLK motif located within the coiled-coil leading up to the RWD domain ^21, 45^; (2) the C-terminus of BUB1 CM1 contacts the RWD domain of the opposite MAD1 ^45^; (3) BUB1 interacts with CDC20 through multiple motifs cooperatively, including the ABBA motif (527–532, which binds between blades 2 and 3 of CDC20’s seven-bladed WD40 fold) and the consensus KEN box (C-terminal to the ABBA motif, which likely binds to the center of CDC20’s WD40) ^16, 46, 47^.

In the parallel study by Fischer and colleagues ^40^, the purified MAD1:MAD2 complex was shown to exhibit a folded conformation *in vitro*. Here, we showed that the MAD1:MAD2 complex may assume such a folded conformation also *in vivo*. Our data indicate that the structural flexibility is enabled by a flexible hinge in the C-terminus of MAD1, whose secondary structure – rather than primary sequence – is conserved. This hinge is important for MCC assembly *in vitro* and SAC signaling *in vivo*, and we provide evidence that it can be replaced with similarly flexible but different sequences, implying that the hinge is unlikely to mediate hitherto unknown physical interactions with other proteins. Thus, collectively, the structural flexibility of MAD1 appears to be important to the SAC signaling activity.

Whether MAD1 switches between an extended conformation and the folded conformation at a physiologically meaningful rate *in vivo*, and whether this switching cycle correlates with the “knitting” of a CDC20:MAD2 heterodimer is currently unclear. The distribution of conformations of the two proline residues (P585 and P596) in the hinge may be under active, energy-consuming regulation in the cell, but assessing this will require further analyses. We note that no MAD1-interacting protein with peptidylprolyl cis-trans isomerase activity has been identified in the PrePPI database as of March 2022 ^41, 42^. It remains unknown whether the proline residues simply serve to break the coiled-coil or play a more complex role in promoting the folding of MAD1.

Our *in vitro* reconstitution data suggest that the critical role of the flexibility of MAD1 is strictly coupled with BUB1. In the absence of BUB1 in the reactions, the assembly rates of CDC20:MAD2 were the same for both MAD1 and MAD1ΔL. However, assembly of MCC, albeit at low rates, continues during interphase and prophase ^43^. There has been no report on BUB1’s localization at the NPC where the MAD1:MAD2 complex is predominantly localized during the interphase and prophase. Therefore, either the flexibility of MAD1 alone scaffolds CDC20:MAD2 coupling at the NPC or there may be a nucleoporin that functions similarly to BUB1. Interestingly, the nuclear basket protein TPR, which is directly associated with the MAD1:MAD2 complex during the interphase and prophase ^44^, is predicted to bind to CDC20 directly in the PrePPI database ^42^. Future studies should look into how the MAD1:MAD2 complex may catalyze the formation of the CDC20:MAD2 dimer at the NPC during the interphase and prophase.

## Author contributions

C.C., A.M., and A.P.J. wrote the manuscript. V.P. contributed to its revision. C.C. performed all human cell experiments. S.J.Y.H. and B.R. performed all experiments related to budding yeasts. V.P. purified recombinant proteins and performed all MCC FRET-sensor-based assays. A.A. performed the pull-down experiment with amylose beads. P.J.H.I.V. performed low-angle metal shadowing and electron microscopy.

## Acknowledgments

This study is funded by NIH grant R35-GM-126983-01 (to A.P.J.). We thank the Single Molecule Analysis in Real-Time Center at the University of Michigan (seeded by NSF MRI-R2-ID award DBI-0959823 to Nils G. Walter) and J. Damon Hoff for technical advice on FLIM. We thank the Flow Cytometry Core at the University of Michigan Medical School for assistance in all flow cytometry experiments. We thank Hongtao Yu (Westlake University, China and University of Texas Southwestern Medical Center, United States) for helpful discussions on the internal tagging of MAD2, as well as Yibo Luo and Song-Tao Liu (University of Toledo, United States) for helpful discussions on mutations of MAD1’s hinge.

## Figure Legends

**Figure S1.**
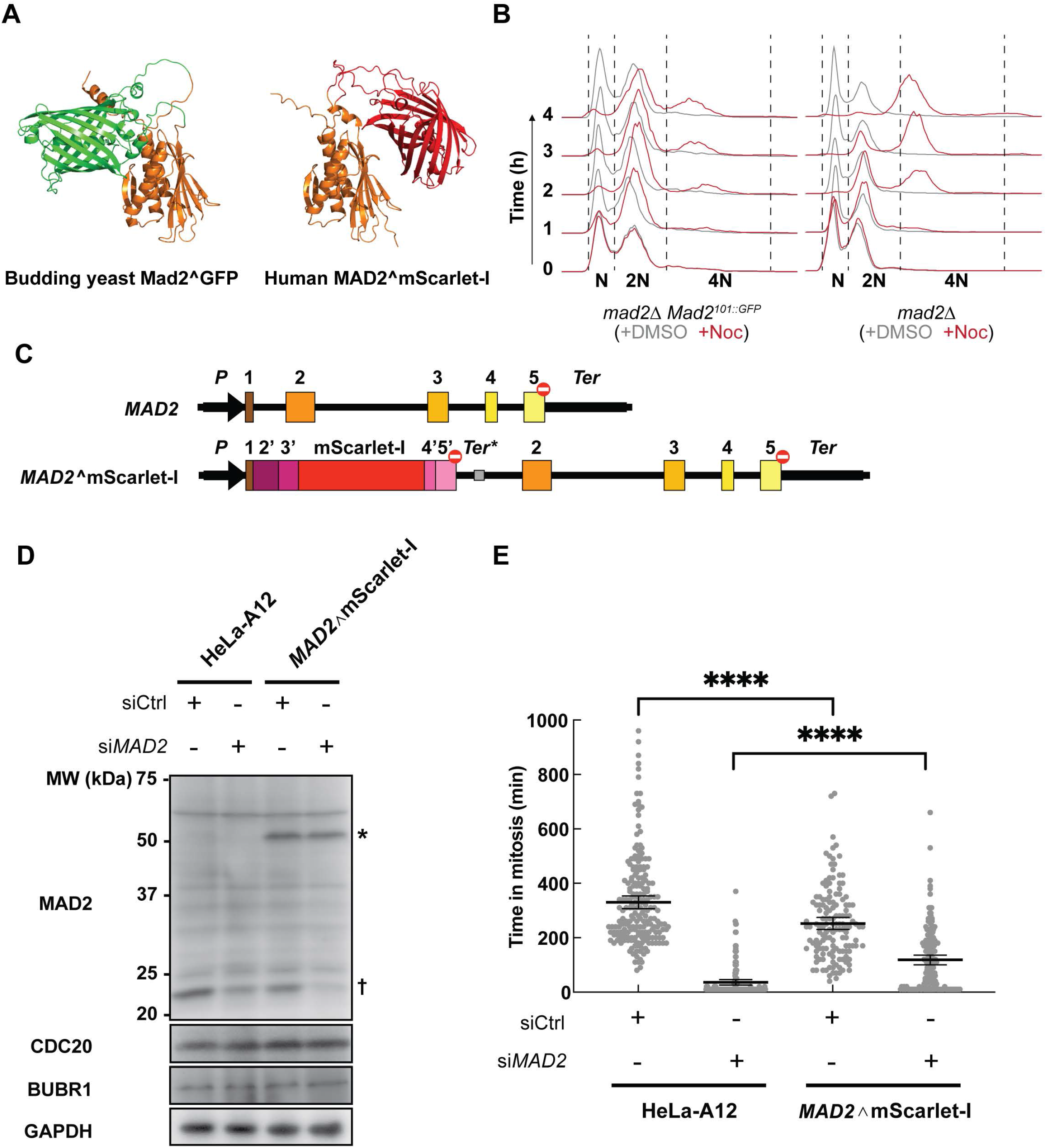
Internally tagged MAD2 is functional in both budding yeast and human cells. (A) A representative model of *Saccharomyces cerevisiae* Mad2∧GFP (left; internally tagged within the β5-αC loop) and human MAD2∧mScarlet-I (right; internally tagged within the β5-αC loop) predicted by the ColabFold advanced algorithm. (B) Effects of Nocodazole treatment on the *mad2Δ S. cerevisiae* strain or the *mad2Δ* strain expressing Mad2∧GFP. The graphs show the quantification of cellular DNA content using flow cytometry 0-4 h after supplementing the growth media with DMSO (gray) or nocodazole (red). Normal interphase cells are haploids whose DNA content corresponds to “N”. Representative results from two experiments were shown. (C) Diagram of the endogenous *MAD2* allele and the genome-edited *MAD2*∧mScarlet-I allele. Boxes 1–5 represent the exons. The regions between these boxes represent the introns. Boxes 2’–5’ encode the same peptides as boxes 2–5 respectively, with the introduction of certain silence mutations that make the exogenous MAD2∧mScarlet-I resistant to siMAD2. The black “*P*” arrow represents the promoter and the 5’-UTR. The black “*Ter*” bar represents the 3’-UTR and the polyadenylation signal. The gray “*Ter*\*” bar represents the polyadenylation signal of rabbit β-globin. The red stop signs represent stop codons. The sequence of the MAD2∧mScarlet-I allele was confirmed by genotyping and Sanger sequencing (data not shown). (D) Immunoblotting showed that MAD2∧mScarlet-I (labeled by an asterisk, with an expected molecular weight of 51.0 kDa) was correctly expressed in the heterozygous *MAD2*∧mScarlet-I HeLa-A12 cell line and was resistant against si*MAD2*. As a comparison, wild-type MAD2 (labeled by a cruciform with a molecular weight of 23.5kDa) was effectively knocked down by si*MAD2*. The immunoblot against GAPDH served as the loading control. (E) Unsynchronized cells were treated with respective siRNAs for one day, treated with 50 nM nocodazole. Each gray dot represents a cell. The total number of cells in each group *N* > 140. Mean values ±95% confidence intervals are overlaid. Results are representative of two independent experiments. Unpaired *t*-tests with Welch’s correction are performed in Prism.

**Figure S2.**
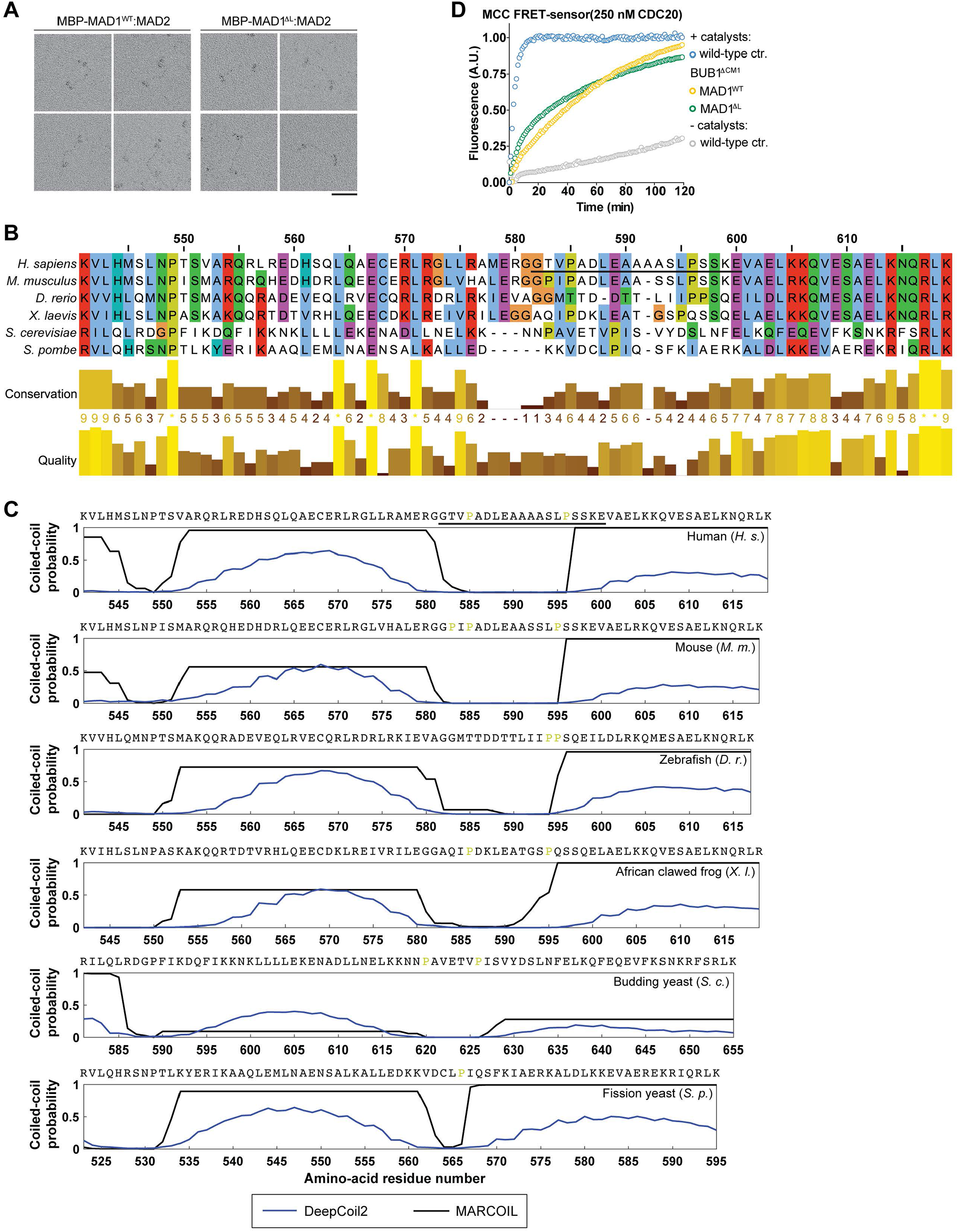
The secondary structure of the hinge of MAD1 is well conserved. (A) MBP-MAD1(wild-type or ΔL):MAD2 visualized by electron microscopy after glycerol spraying and low-angle platinum shadowing. Scale bars, 50 nm. (B) The primary sequence of MAD1’s hinge is not conserved. Jalview is used in the multiple sequence alignment (using the MSAprobs alignment tool with default settings) and visualization (Waterhouse et al. 2009) and the coloring scheme of Clustal X is applied. The amino-acid residue numbering at the top is for human MAD1. (C) The presence of this flexible hinge in the C-terminus of MAD1 is conserved and proline residues (colored yellow) are usually present within this region. The figure shows coiled-coil predictions by two algorithms (blue curves: raw predicted probabilities by DeepCoil2; black curves: MARCOIL) on the region spanning from MAD1’s MIM (which is also not a coiled-coil ^13^) to MAD1’s consensus RLK motif from *Homo sapiens* (human), *Mus musculus* (mouse), *Danio rerio* (zebrafish), *Xenopus Laevis* (African clawed frog), *Saccharomyces cerevisiae* (budding yeast), and *Schizosaccharomyces pombe* (fission yeast). The RLK motif directly binds to BUB1 ^21, 45^ and is located within the coiled-coil leading to the RWD domain (see the crystal structure on the right in Figure 1A). The primary sequences of full-length MAD1 proteins were supplied as the input, but only probability predictions for the region spanning from the MIM to the RLK motif are shown. Similar prediction results were obtained using PSIPRED 4.0 ^48, 49^, although the exact starting and ending residues of the flexible hinge may differ (data not shown). In both (B) and (C), the segment encompassing residues 582–600 of human MAD1 is underlined. (D) MCC FRET-sensor-based assays show that when BUB1ΔCM1 is used instead of wild-type BUB1, MBP-MAD1:MAD2 (yellow) and MBP-MAD1ΔL:MAD2 (green) have a similar decreased activity in promoting MCC assembly. Curves report single measurements representative of at least three independent technical replicates. Prism was used for data analysis and visualization.

**Figure S3.**
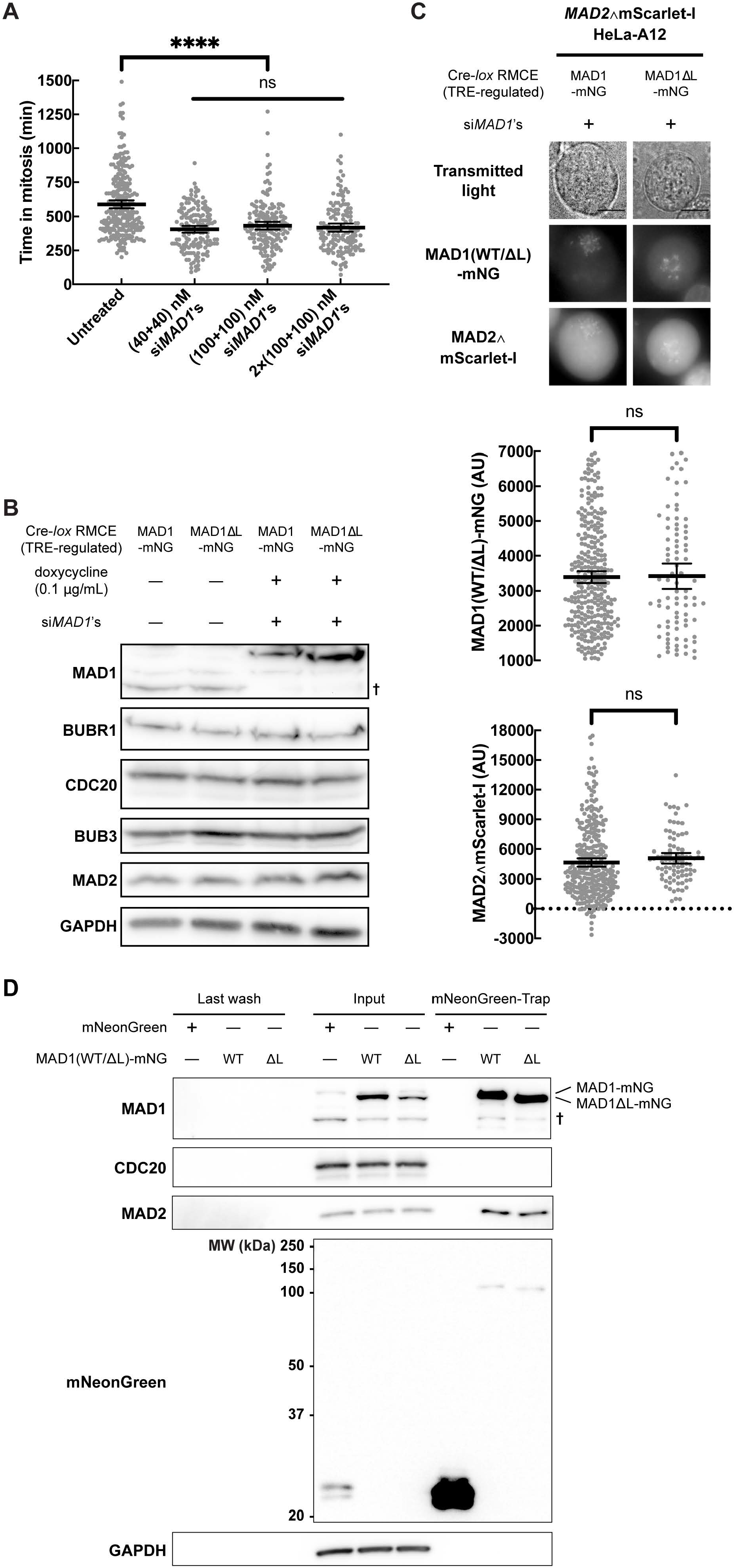
Deletion of the hinge does not affect the localization of the MAD1:MAD2 complex or the expression level of MCC constituents. (A) MAD1 has a long half-life under normal conditions ^50^. And like BUB1 ^51-53^, even a small pool of MAD1 (at less than 10% of its physiological concentration as quantified from Figure S3B) can maintain a considerable level of SAC signaling activity in nocodazole-treated cells. The conditions of si*MAD1* treatment were (from left to right): untreated, 40nM each for two days (the standard condition used throughout this study), 100 nM each for two days, 100 nM each on day one and 100 nM each again on day two. The *MAD1*-mNG genome-edited HeLa-A12 cell line was used in each group. Each dot represents a cell (*N* ≥ 145 in each group). The mean value ± the 95% confidence interval of each group is overlaid. Welch’s ANOVA test [*W*(DF*n*, DF*d*) = 0.9885(2.000, 298.9), *p* = 0.3733] was performed for the three columns on the right. The ANOVA test and the unpaired *t*-test with Welch’s correction are performed in Prism. (B) Knockdown of the endogenous *MAD1* by si*MAD1*’s had an efficiency of over 90% based on the intensity of the residual MAD1 band. The cellular abundance of either BUBR1, CDC20, or BUB3 was not affected. The immunoblot against GAPDH served as the loading control. (C) The *MAD2*∧mScarlet-I genome-edited HeLa-A12 treated with si*MAD1*’s and rescued by MAD1(WT/ΔL)-mNG were imaged using wide-field fluorescence microscopy. Cells were arrested at mitosis using a thymidine–nocodazole synchronization protocol. Representative micrographs are shown in the top panel. Maximum *z*-projected green channel images shown here share the same LUT. Maximum *z*-projected red channel images shown here also share the same LUT. Scale bar, 10 μm. Due to various expression levels of induced MAD1(WT/ΔL)-mNG in different cells, signaling kinetochores were filtered by the localization of MAD1(WT/ΔL)-mNG (with an arbitrary threshold of 1000–7000AU). Each gray dot represents a single signaling kinetochore (*N* ≥ 85 in each group). The mean value ± the 95% confidence interval of each group is overlaid. Unpaired *t*-tests with Welch’s correction are performed in Prism. (D) Using immunoblotting to evaluate the immunoprecipitation by the mNeonGreen-Trap Agarose. The cruciform symbol represents the endogenous MAD1 band. The expected molecular weights of the exogenous MAD1-mNG, MAD1ΔL-mNG, and mNeonGreen are 110.2 kDa, 108.4 kDa, and 26.9 kDa, respectively. The immunoblot against GAPDH served as the loading control. The immunoblots shown here are from the same immunoprecipitation experiment representative of two independent repeats.

**Figure S4.**
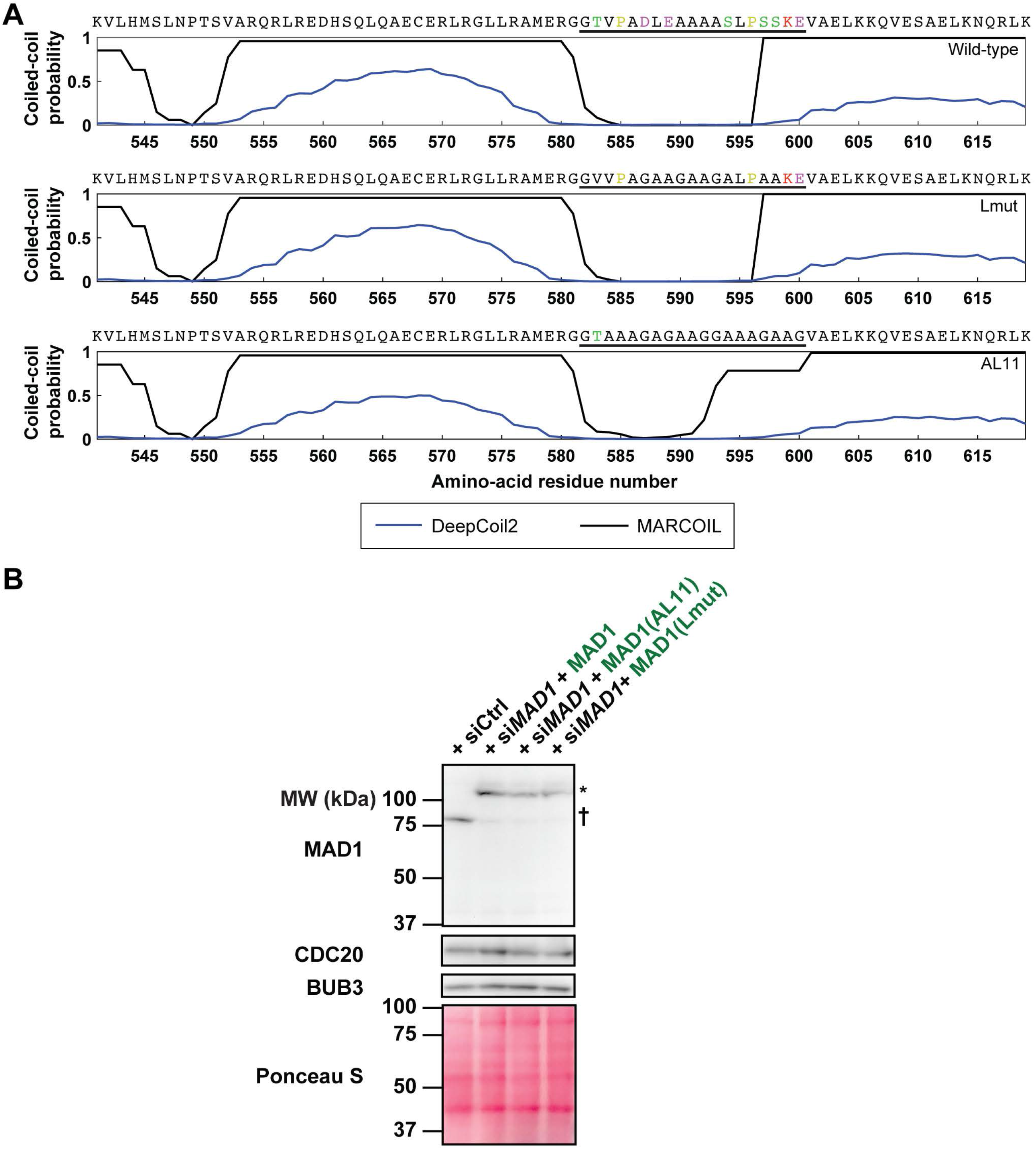
Coiled-coil predictions of MAD1(Lmut/AL11) reveal similar propensity profile as wild-type MAD1. (A) The two algorithms and legends are the same as in Figure S2C. The top panel is reproduced from Figure S2C. Segments encompassing residues 582–600 are underlined. Serine/threonine residues are colored green. Proline residues are colored yellow. Negatively charged residues are colored purple. Lysine residues are colored red. (B) Immnunoblot analysis of mitotic lysates of HeLa-A12 cells (first lane from the left) or HeLa-A12 cells expressing exogenous MAD1-mNG (second lane), MAD1(AL11)-mNG (third lane), or MAD1(Lmut)-mNG (fourth lane). Cells were treated with labeled siRNAs and 0.1 μg/mL doxycycline for two days. Cells were synchronized by 2.5 mM thymidine overnight, released for 7 h, and treated with 330nM nocodazole for 4h before being harvested by the mitotic shake-off technique. The expected molecular weights of the exogenous MAD1-mNG, MAD1(AL11)-mNG, and MAD1(Lmut)-mNG are 110.2 kDa, 109.7 kDa, and 110.0 kDa, respectively. Ponceau S staining (bottome panel) of the MAD1 blot serves as a control for sample loading and membrane transfer.

## Materials and methods

For methods of cell culture and Cre-*lox* RMCE, see [1]. Wide-field, *z*-stack fluorescence imaging used in the quantification of localization of MAD1(WT/ΔL)-mNG and MAD2∧mScarlet-I at signaling kinetochores was the same as described in [2]. AlphaFold 2 structure predictions were conducted using the ColabFold advanced algorithm. All the parameters were set at their default values except for “max recycles” (which was set to 6) and “tol” (which was set to 0.1).

### Theoretical end-to-end root-mean-square distance of a flexible unstructured peptide

First, we model a flexible peptide with *n* amino acid residues using a 3-D random walk model (without considering steric hindrance and restrictions imposed by the Ramachandran plot). We denote the displacement of residue number *i*+1 relative to residue number *i* as a random vector **r**_*i*_, *i* = 1, 2, …, *n* 1. The end-to-end displacement, **D**, can be expressed as

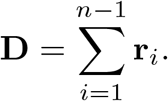

The root mean square of it is therefore

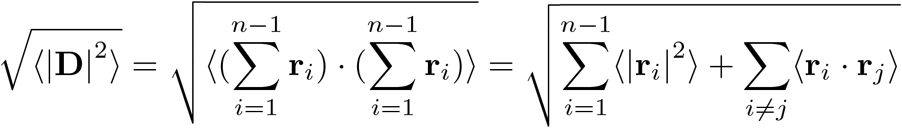

For a 3-D random walk, the random vectors representing each step are independent of each other. Therefore, ∀*i* ≠ *j*,

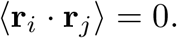

Suppose that the contour length of each amino acid residue is universal (|**r**_*i*_| = *r, i* = 1, 2, …, *n* − 1; according to [3], we take *r* = 0.37nm here), we have

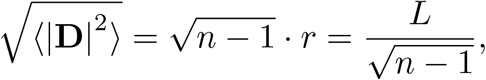

wherein *L* = (*n* − 1)*r* is the contour length of the peptide.

Second, we model the same peptide using a worm-like chain model. This model considers the peptide as a continuous worm-like chain rather than a discrete, step-by-step walk in the previous model. According to [4], the end-to-end root-mean-square distance

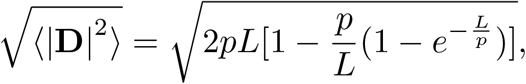

wherein *p* is the persistence length, a metric for the stiffness of the chain. According to [4, 5], we take *p* = 0.3–0.7nm here.

### Purification of recombinant proteins

Wild-type or mutant constructs of MAD1:MAD2, MAD2, MPS1, BUB1:BUB3, CDC20, and BUBR1: BUB3 are of human origin. The constructs of MBP-MAD1ΔL:MAD2 and MBP-MAD1(AL11):MAD2 are cloned via site-directed mutagenesis from the MBP-MAD1:MAD2 wild-type construct used in [6, 7]. All recombinant proteins used in this study have been expressed and purified according to the protocols described in [6, 7].

### Low-angle metal shadowing and electron microscopy

MBP-MAD1(wild-type or ΔL):MAD2 was diluted 1 : 1 with a spraying buffer (200mM ammonium acetate and 60% glycerol) to a final concentration of 0.5–1.0 µM and air-sprayed onto freshly cleaved mica pieces (V1 quality, Plano GmbH). Specimens were mounted and dried in a MED020 high-vacuum metal coater (Bal-tec). A platinum layer of approximately 1nm and a 7-nm carbon support layer were subsequently evaporated onto the rotating specimen at angles of 6–7° and 45°, respectively. Pt/C replicas were released from the mica on water, captured by freshly glow-discharged 400-mesh Pd/Cu grids (Plano GmbH), and visualized using a LaB6 equipped JEM-1400 transmission electron microscope (JEOL) operated at 120 kV. Images were recorded at a nominal magnification of 60, 000× on a 4k×4k CCD camera F416 (TVIPS).

### FRET assay with the MCC FRET-sensor

The MCC FRET-sensor has been described previously [6, 7]. The catalysts preparation consisted of 2 µM MBP-MAD1(wild-type or mutant):MAD2 and 2 µM BUB1(wild-type or mutant):BUB3, which were separately incubated with 500nM MPS1 in the assay buffer [10mM HEPES (pH 7.5), 150mM NaCl, 2.5% glycerol, and 10mM β-mercaptoenthanol] supplemented with 1mM ATP and 10mM MgCl_2_ for 16 h at 4 °C. All assays were performed using 100nM final concentration of all proteins, except for CDC20, which was added at 250nM (instead of 500nM used in previous studies [6, 7]). The fluorophores MAD2-TAMRA and mTurquoise-BUBR1(1-571):BUB3 were added before measurements started. All measurements were performed on a CLARIOstar plate reader (BMG Labtech), using UV-Star 96-well plates (Greiner). The reactions had a final volume of 100 µl in the assay buffer. The excitation light and emitted fluorescence were filtered by a 430-10 nm excitation filter, an LP 504 nm dichroic mirror, and a 590-20 nm emission filter. The plate reader read at a 60 s interval for 120 min (6mm focal height, 200 flashes, gain 1200) and mix the reactions for 5 s at 500 rpm after each measurement.

### Flow cytometry

The complete genotype of the *mad2Δ S. cerevisiae* strain (AJY4951, [8]) is *leu2Δ-1, trp1Δ63, ura3-52, his3Δ200, lys2-8Δ1, mad2Δ::TRP1*. The complete genotype of the Mad2∧GFP-expressing *S. cerevisiae* strain (AJY5041, constructed in this study) is *leu2Δ0, met15Δ0, ura3Δ0, mad2Δ::KAN, Mad2*^*101::GFP*^ *(HIS3)*. Yeast strains were grown to mid-log phase and then 15 µg*/*mL nocodazole was added to the media. Sample aliquots containing ∼ 2 × 10^6^ cells were collected 0, 1, 2, 3, and 4 h after the addition of nocodazole. Samples were fixed by 70% ethanol and then stored at 4 °C overnight. On day two, samples were washed and treated with 170 ng*/*mL bovine pancreatic RNase (Millipore Sigma) at 37 °C for one day in the RNase buffer [10mM Tris (pH 8.0) and 15mM NaCl]. On day three, samples were washed again, resuspended in PBS, and stored at 4 °C. The samples were treated with 5 mg*/*ml propidium iodide (Millipore Sigma) for 2 h at room temperature and subject to flow cytometry on an LSRFortessa Cell Analyzer (BD Biosciences). Data were analyzed using FlowJo.

### Generating the *MAD2*∧mScarlet-I genome-edited HeLa-A12 cell line

The gRNA used in the integration of the coding sequence of MAD2∧mScarlet-I (intron-free, stop codon-containing, and si*MAD2*-resistant by the introduction of silent mutations) and the polyadenylation signal of rabbit β-globin after the first exon of the endogenous *MAD2* gene was 5’-UCGCG CAGGCCAAUAUAUCG-3’. Synthesis of the sgRNA and assembly of the *Sp*Cas9-sgRNA RNP complex were described in [9]. Plain or *MAD1*-mNG genome-edited HeLa-A12 cell lines were co-transfected with the RNP complex and linearized pCC35, sorted, and validated as described in [9]. A successfully edited *MAD2*∧mScarlet-I allele encodes an internally-tagged MAD2 protein, wherein wild-type MAD2 and mScarlet-I are separated by short flexible linkers (`AGSGSGGAS` between S114 of MAD2 and the N-terminus of mScarlet-I; `GTGAGSA` between the C-terminus of mScarlet-I and A115 of MAD2).

### RNA interference

The two siRNAs targeting the 3’-UTR of *MAD1* were from [10]. They were applied to unsynchronized cells at a concentration of 40nM each for two days before imaging or collecting cells for immunoblotting unless specified otherwise. The sense-strand sequence of si*CDC20* was 5’-GGAGCUCAUCUCAGGCCAU-3’ [11], which was applied at a concentration of 40nM for two days before FLIM or immunoblotting. The sense-strand sequence of si*MAD2* was 5’-GGAAGAGUCGGGACCACAGUU-3’ [12], which was applied at a concentration of 40nM for one day before imaging or immunoblotting. Desalted siRNAs modified by double-deoxythymidine overhangs at 3’-ends of both strands were synthesized by Sigma. AllStars Negative Control siRNA (QIAGEN) is used as the control siRNA (siCtrl) and applied at the same dosage and time as the corresponding experimental group(s). All siRNAs were transfected into the cells via Lipofectamine RNAiMAX following manufacturer’s instructions.

### Fluorescence lifetime imaging microscopy (FLIM)

All FLIM data were collected on an Alba v5 Laser Scanning Microscope, connected to an Olympus IX81 inverted microscope main body [equipped with a UPLSAPO60XW objective (1.2 NA)]. A Fianium WL-SC-400-8 laser with an acousto-optic tunable filter was used to generate excitation pulses at a wavelength of 488nm and a frequency of about 20 MHz. Excitation light was further filtered by a Z405/488/561/635rpc quadband dichroic mirror. The emission light of the green channel was redirected by a 562 longpass dichroic mirror (FF562-Di03, Semrock), filtered by an FF01-531/40-25 filter, and finally detected by an SPCM-AQRH-15 avalanche photodiode. The time-correlated single photon counting module to register detected photon events to excitation pulses was SPC-830. Data acquisition was facilitated by VistaVision.

The emission light was redirected by a 562 longpass dichroic mirror and filtered by an FF01-582/75-25 filter (Semrock). The data analysis pipeline (implemented in MATLAB) developed in this study is publicly available on https://github.com/CreLox/FluorescenceLifetime.

To demonstrate how fluorescence lifetime measurements can quantify the FRET efficiency, consider donor fluorophores with a lifetime of *τ*_0_. In the absence of acceptor fluorophores, the exponential decay *D*_0_ of donor fluorescence after the pulse excitation at time zero is

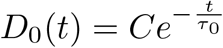

and the total fluorescence signal is

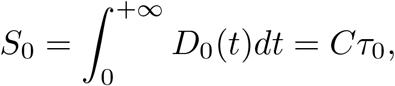

wherein *C* is a constant determined by the excitation and detection condition, the total number and properties of fluorophores, and the imaging setup. Without altering any of these conditions, in the presence of acceptor fluorophores and FRET, the longer a donor fluorophore stays excited, the higher the chance FRET may have occurred (note: this is not a rigorous statement because fluorescence emission and FRET quenching are independent stochastic processes and an excited fluorophore can only relax through one route). Suppose that the timing of FRET follows an exponential distribution with a probability density function of

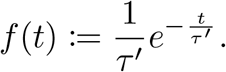

The probability that FRET does not happen before *t*_0_ will be

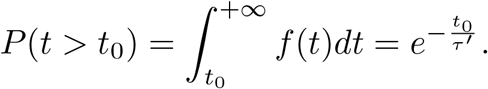

Excited fluorophores can either take the FRET quenching route or the fluorescence emission route to relax to the ground state (note: a fluorophore may also relax through other ways but the fact that these routes are independent stochastic processes means that it does not affect the following conclusion). Therefore, in the presence of acceptor fluorophores and FRET, the decay *D* of donor fluorescence becomes

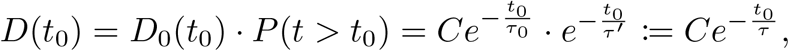

wherein the new lifetime is

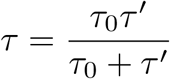

and the new total fluorescence signal is *S* = *Cτ*. Therefore,

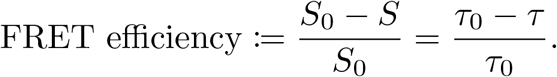

Because the fluorescence lifetime in the absence of quenching is an intrinsic property of a mature fluorescent protein (under a certain temperature) [13], the equation above greatly simplifies experiments to measure the FRET efficiency. This equation still applies even if the fluorescence decay has to be fitted by a multi-component exponential decay, as long as the fluorescence lifetime is an average weighted by the corresponding *C* of each component.

### Time-lapse live-cell imaging in knockdown-rescue mitotic duration assays

Time-lapse live-cell imaging was performed on an ImageXpress Nano Automated Imaging System (Molecular Devices). A SOLA Light Engine (Lumencor) served as the excitation light source. Cells were plated on 24-well cell imaging plates (black plate with treated glass bottom, Eppendorf) and treated with siRNAs and 100nM nocodazole accordingly. Humidified 5% CO_2_ was supplied to the environment chamber maintained at 37 °C.

According to [14], the level of MAD1 and MAD2 has to be balanced for a robust SAC. To make sure that the expression of exogenous, si*MAD1*-resistant MAD1(wild-type/mutant)-mNG in si*MAD1*-treated cells is close to the physiological level of endogenous MAD1 for all analyzed cells, we image the heterozygous *MAD1*-mNG genome-edited HeLa-A12 cell line [9] as the control in all of our knockdown-rescue mitotic duration assays. In this cell line, the stoichiometry of endogenous mNG-tagged versus untagged MAD1 is about 1 : 1 [9]. Therefore, only cells with mNG intensity (after background and shading correction) close to two times the mNG intensity in the heterozygous *MAD1*-mNG genome-edited HeLa-A12 cell line were analyzed in our knockdown-rescue mitotic duration assays (Figures 3A and 3C).

### Pull-down using amylose beads

BUB1:BUB3, CDC20, O-MAD2(V193N), and MBP-MAD1(wild-type or mutant):MAD2 were diluted using a binding buffer [20mM HEPES (pH 7.5), 150mM NaCl, 5% glycerol] in a total volume of 50 µL. Unless specified otherwise, MBP-MAD1(wild-type or mutant):MAD2 and BUB1:BUB3 were diluted at 20 µM and pre-phosphorylated at 4 °C for 16 h by MPS1 (1 µM). The final concentration of MBP, MBP-MAD1(wild-type or mutant):MAD2 was 4 µM; the final concentration of BUB1:BUB3, CDC20, and O-MAD2(V193N) were 5 µM each. 50 µL of the solution was mixed with 15 µL of amylose beads (New England Biolabs). Samples were placed into PierceTM micro-spin columns (Thermofischer) and incubated at 4 °C for 1 h. To separate the proteins bound to the amylose beads from the unbound proteins, the samples were centrifuged at 900 g for 2 min at 4 °C. The beads were washed three times with 200 µL of binding buffer. After the last washing step, 25 µL of elution buffer (binding buffer plus 10 mM maltose) was added to the column and centrifuged at 800 g for 2 min at 4 °C. The eluted proteins were mixed with 5× SDS-PAGE loading buffer and analyzed by SDS-PAGE and immunoblotting.

### Immunoprecipitation using mNeonGreen-Trap

HeLa-A12 cells integrated with the Tet-On expression cassette of either mNeonGreen, MAD1-mNG, or MAD1ΔL-mNG were induced to express the ectopic exogenous protein by 0.1 µg*/*mL doxycycline (for two days until being harvested) and arrested at mitosis using a thymidine–nocodazole synchronization protocol. Cells were harvested by mitotic shake-off, washed once by PBS, pelleted down by centrifugation at 200–500 g for 3 min, snap-frozen in liquid nitrogen, and stored at −80 °C before the immunoprecipitation (IP) experiment.

On the day of the immunoprecipitation experiment, cells were thawed on ice and lysed in the IP lysis buffer [75mM HEPES-HCl (pH 7.5 at 4 °C), 150mM KCl, 10% (by volume) glycerol, 1.5mM MgCl_2_, 1.5mM EGTA, and 1% (by mass) CHAPS, a zwitterionic detergent] supplemented before usage with 1mM PMSF, the cOmplete™ EDTA-free Protease Inhibitor Cocktail, Phosphatase Inhibitor Cocktail IV (RPI), and a phosphatase inhibitor cocktail (1mM Na4P2O7, 0.1mM Na3VO4, 5mM NaF, and 2mM sodium β-glycerophosphate). For 1 mg of wet cell pellet, 40 µL of 4 °C IP lysis buffer was added, yielding a total protein concentration of about 5.6 mg*/*mL (if cells were lysed completely). Resuspended cells were rotated for 30 min at 4 °C and then centrifuged at 18,000 g for 20 min at 4 °C. 600 µL of supernatant was subsequently cleared by 50 µL of equilibrated control agarose beads (ChromoTek) to reduce non-specific bindings, rotating for 45 min at 4 °C. The mixture was centrifuged at 2000 g for 5 min at 4 °C. 580 µL of pre-cleared supernatant was then mixed with 30 µL of equilibrated mNeonGreen-Trap Agarose (nta-20, ChromoTek) and rotated for 1 h at 4 °C. These beads were then pelleted down at 2000 g for 5 min at 4 °C and the supernatant was removed. The beads were further washed four times (rotated for 5 min at 4 °C and then pelleted down at 2000 g for 5 min at 4 °C) using 1 mL of the IP wash buffer [75mM HEPES-HCl (pH 7.5 at 4 °C), 150mM KCl, 10% (by volume) glycerol, 1.5mM MgCl_2_, and 1.5mM EGTA] each time. The beads were transferred to a fresh tube before the last wash to avoid the non-specific binding of proteins to the wall of the tube. Finally, 2× Laemmli buffer supplemented with β-mercaptoethanol was added to the beads. Samples were boiled in a boiling water bath for 10 min before being subjected to SDS-PAGE and immunoblotting analysis.

### Immunoblotting

To acquire unsynchronized HeLa-A12 cells, asynchronous cells were either scrapped or trypsinized off the surface of dishes. To acquire mitotic HeLa-A12 cells, cells were first synchronized in G1/S with 2.5mM thymidine and then arrested in mitosis with 330nM of nocodazole for 16 h. This procedure is referred to as the thymidine–nocodazole synchronization protocol in the main text.

Harvested cells were then washed once by PBS, pelleted down, and chilled on ice. Lysis was performed by directly adding 2× Laemmli sample buffer (Bio-Rad Laboratories, supplemented by 2-mercaptoethanol) at a ratio of 1 µL per 0.1 mg of cell pellets and pipetting up and down. Lysates were boiled immediately afterward for 10 min and then chilled on ice. 8 µL of supernatant was loaded onto each lane of a 15-well, 0.75-mm SDS-PAGE mini gel.

Primary antibodies (and their working dilution factors by volume) used included anti-BUBR1 (Bethyl Laboratories A300-995A-M, 1 : 1000), anti-BUB1 (Abcam ab9000), anti-CDC20 (Santa Cruz Biotechnology sc-5296 for Figure 2F and sc-13162, 1 : 200 for others), anti-MAD2 (Bethyl Laboratories A300-301A-M, 1 : 330), anti-GAPDH (Proteintech 60004-1-Ig, 1 : 5000), anti-MAD1 (GeneTex GTX109519, 1 : 2000 in Figure 1E and PLA0092, 1 : 1000 in Figures S3B and S3D), anti-mNeonGreen (Cell Signaling Technology 53061S, 1 : 100), and anti-BUB3 (Sigma-Aldrich B7811, 1 : 500).

## References

1. Sudakin, V., Chan, G.K. & Yen, T.J. Checkpoint inhibition of the APC/C in HeLa cells is mediated by a complex of BUBR1, BUB3, CDC20, and MAD2. J Cell Biol 154, 925–936 (2001).

2. Chao, W.C., Kulkarni, K., Zhang, Z., Kong, E.H. & Barford, D. Structure of the mitotic checkpoint complex. Nature 484, 208–213 (2012).

3. Sudakin, V. et al. The cyclosome, a large complex containing cyclin-selective ubiquitin ligase activity, targets cyclins for destruction at the end of mitosis. Mol Biol Cell 6, 185–197 (1995).

4. Alfieri, C. et al. Molecular basis of APC/C regulation by the spindle assembly checkpoint. Nature 536, 431–436 (2016).

5. Yamaguchi, M. et al. Cryo-EM of Mitotic Checkpoint Complex-Bound APC/C Reveals Reciprocal and Conformational Regulation of Ubiquitin Ligation. Mol Cell 63, 593–607 (2016).

6. Clute, P. & Pines, J. Temporal and spatial control of cyclin B1 destruction in metaphase. Nat Cell Biol 1, 82–87 (1999).

7. Yu, J. et al. Structural basis of human separase regulation by securin and CDK1-cyclin B1. Nature 596, 138–142 (2021).

8. Chang, D.C., Xu, N. & Luo, K.Q. Degradation of cyclin B is required for the onset of anaphase in Mammalian cells. J Biol Chem 278, 37865–37873 (2003).

9. Simonetta, M. et al. The influence of catalysis on mad2 activation dynamics. PLoS Biol 7, e10 (2009).

10. Faesen, A.C. et al. Basis of catalytic assembly of the mitotic checkpoint complex. Nature (2017).

11. Luo, X., Tang, Z., Rizo, J. & Yu, H. The Mad2 spindle checkpoint protein undergoes similar major conformational changes upon binding to either Mad1 or Cdc20. Mol Cell 9, 59–71 (2002).

12. Luo, X. et al. Structure of the Mad2 spindle assembly checkpoint protein and its interaction with Cdc20. Nat Struct Biol 7, 224–229 (2000).

13. Sironi, L. et al. Crystal structure of the tetrameric Mad1-Mad2 core complex: implications of a ‘safety belt’ binding mechanism for the spindle checkpoint. EMBO J 21, 2496–2506 (2002).

14. De Antoni, A. et al. The Mad1/Mad2 complex as a template for Mad2 activation in the spindle assembly checkpoint. Curr Biol 15, 214–225 (2005).

15. Luo, X. et al. The Mad2 spindle checkpoint protein has two distinct natively folded states. Nat Struct Mol Biol 11, 338–345 (2004).

16. Piano, V. et al. CDC20 assists its catalytic incorporation in the mitotic checkpoint complex. Science 371, 67–71 (2021).

17. Lara-Gonzalez, P., Kim, T., Oegema, K., Corbett, K. & Desai, A. A tripartite mechanism catalyzes Mad2-Cdc20 assembly at unattached kinetochores. Science 371, 64–67 (2021).

18. Kim, S., Sun, H., Tomchick, D.R., Yu, H. & Luo, X. Structure of human Mad1 C-terminal domain reveals its involvement in kinetochore targeting. Proceedings of the National Academy of Sciences 109, 6549–6554 (2012).

19. Kruse, T. et al. A direct role of Mad1 in the spindle assembly checkpoint beyond Mad2 kinetochore recruitment. EMBO reports 15, 282–290 (2014).

20. Heinrich, S. et al. Mad1 contribution to spindle assembly checkpoint signalling goes beyond presenting Mad2 at kinetochores. EMBO Rep 15, 291–298 (2014).

21. Ji, Z., Gao, H., Jia, L., Li, B. & Yu, H. A sequential multi-target Mps1 phosphorylation cascade promotes spindle checkpoint signaling. Elife 6 (2017).

22. Ji, W., Luo, Y., Ahmad, E. & Liu, S.T. Direct interactions of mitotic arrest deficient 1 (MAD1) domains with each other and MAD2 conformers are required for mitotic checkpoint signaling. J Biol Chem 293, 484–496 (2018).

23. Zhou, H.X. Polymer models of protein stability, folding, and interactions. Biochemistry 43, 2141–2154 (2004).

24. Lapidus, L.J., Steinbach, P.J., Eaton, W.A., Szabo, A. & Hofrichter, J. Effects of Chain Stiffness on the Dynamics of Loop Formation in Polypeptides. Appendix: Testing a 1-Dimensional Diffusion Model for Peptide Dynamics. The Journal of Physical Chemistry B 106, 11628–11640 (2002).

25. Jumper, J. et al. Highly accurate protein structure prediction with AlphaFold. Nature 596, 583–589 (2021).

26. Mirdita, M., Steinegger, M., Breitwieser, F., Soding, J. & Levy Karin, E. Fast and sensitive taxonomic assignment to metagenomic contigs. Bioinformatics (2021).

27. Bindels, D.S. et al. mScarlet: a bright monomeric red fluorescent protein for cellular imaging. Nat Methods 14, 53–56 (2017).

28. Mapelli, M., Massimiliano, L., Santaguida, S. & Musacchio, A. The Mad2 conformational dimer: structure and implications for the spindle assembly checkpoint. Cell 131, 730–743 (2007).

29. Hara, M., Ozkan, E., Sun, H., Yu, H. & Luo, X. Structure of an intermediate conformer of the spindle checkpoint protein Mad2. Proc Natl Acad Sci USA 112, 11252–11257 (2015).

30. Kukreja, A.A., Kavuri, S. & Joglekar, A.P. Microtubule Attachment and Centromeric Tension Shape the Protein Architecture of the Human Kinetochore. Curr Biol 30, 4869–4881 e4865 (2020).

31. Becker, W. Fluorescence lifetime imaging--techniques and applications. J. Microsc. 247, 119–136 (2012).

32. Ludwiczak, J., Winski, A., Szczepaniak, K., Alva, V. & Dunin-Horkawicz, S. DeepCoil-a fast and accurate prediction of coiled-coil domains in protein sequences. Bioinformatics 35, 2790–2795 (2019).

33. Delorenzi, M. & Speed, T. An HMM model for coiled-coil domains and a comparison with PSSM-based predictions. Bioinformatics 18, 617–625 (2002).

34. Zimmermann, L. et al. A Completely Reimplemented MPI Bioinformatics Toolkit with a New HHpred Server at its Core. J Mol Biol 430, 2237–2243 (2018).

35. Khandelia, P., Yap, K. & Makeyev, E.V. Streamlined platform for short hairpin RNA interference and transgenesis in cultured mammalian cells. Proc Natl Acad Sci USA 108, 12799–12804 (2011).

36. Ballister, E.R., Riegman, M. & Lampson, M.A. Recruitment of Mad1 to metaphase kinetochores is sufficient to reactivate the mitotic checkpoint. J Cell Biol 204, 901–908 (2014).

37. Chen, C. et al. Ectopic Activation of the Spindle Assembly Checkpoint Signaling Cascade Reveals Its Biochemical Design. Curr Biol 29, 104–119 e110 (2019).

38. Alfonso-Perez, T., Hayward, D., Holder, J., Gruneberg, U. & Barr, F.A. MAD1-dependent recruitment of CDK1-CCNB1 to kinetochores promotes spindle checkpoint signaling. J Cell Biol 218, 1108–1117 (2019).

39. Robinson, C.R. & Sauer, R.T. Optimizing the stability of single-chain proteins by linker length and composition mutagenesis. Proc Natl Acad Sci USA 95, 5929–5934 (1998).

40. Fischer, E.S. et al. Juxtaposition of Bub1 and Cdc20 on phosphorylated Mad1 during catalytic mitotic checkpoint complex assembly. bioRxiv, 2022.2005.2016.492081 (2022).

41. Mi, H. et al. Protocol Update for large-scale genome and gene function analysis with the PANTHER classification system (v.14.0). Nat Protoc 14, 703–721 (2019).

42. Zhang, Q.C., Petrey, D., Garzon, J.I., Deng, L. & Honig, B. PrePPI: a structure-informed database of protein-protein interactions. Nucleic Acids Res 41, D828–833 (2013).

43. Rodriguez-Bravo, V. et al. Nuclear pores protect genome integrity by assembling a premitotic and Mad1-dependent anaphase inhibitor. Cell 156, 1017–1031 (2014).

44. Lee, S.H., Sterling, H., Burlingame, A. & McCormick, F. Tpr directly binds to Mad1 and Mad2 and is important for the Mad1-Mad2-mediated mitotic spindle checkpoint. Genes Dev 22, 2926–2931 (2008).

45. Fischer, E.S. et al. Molecular mechanism of Mad1 kinetochore targeting by phosphorylated Bub1. EMBO Rep, e52242 (2021).

46. Tian, W. et al. Structural analysis of human Cdc20 supports multisite degron recognition by APC/C. Proc Natl Acad Sci USA 109, 18419–18424 (2012).

47. Di Fiore, B. et al. The ABBA Motif Binds APC/C Activators and Is Shared by APC/C Substrates and Regulators. Developmental Cell 32, 358–372 (2015).

48. Jones, D.T. Protein secondary structure prediction based on position-specific scoring matrices. J Mol Biol 292, 195–202 (1999).

49. Buchan, D.W.A. & Jones, D.T. The PSIPRED Protein Analysis Workbench: 20 years on. Nucleic Acids Res 47, W402–W407 (2019).

50. Schweizer, N. et al. Spindle assembly checkpoint robustness requires Tpr-mediated regulation of Mad1/Mad2 proteostasis. J Cell Biol 203, 883–893 (2013).

51. Raaijmakers, J.A. et al. BUB1 Is Essential for the Viability of Human Cells in which the Spindle Assembly Checkpoint Is Compromised. Cell Rep 22, 1424–1438 (2018).

52. Rodriguez-Rodriguez, J.A. et al. Distinct Roles of RZZ and Bub1-KNL1 in Mitotic Checkpoint Signaling and Kinetochore Expansion. Curr Biol 28, 3422–3429 e3425 (2018).

53. Zhang, G. et al. Efficient mitotic checkpoint signaling depends on integrated activities of Bub1 and the RZZ complex. EMBO J 38 (2019).

## References

1. Chen, C. et al. Ectopic activation of the spindle assembly checkpoint signaling cascade reveals its biochemical design. Current Biology 29, 104–119 (2019).

2. Chen, C., Humphrey, L., Jema, S., Ferrari, F. & Joglekar, A. P. Bub1 availability limits the Spindle Assembly Checkpoint signaling strength of human kinetochores. bioRxiv (2022).

3. Ainavarapu, S. R. K. et al. Contour length and refolding rate of a small protein controlled by engineered disulfide bonds. Biophysical Journal 92, 225–233 (2007).

4. Zhou, H.-X. Polymer models of protein stability, folding, and interactions. Biochemistry 43, 2141–2154 (2004).

5. Lapidus, L. J., Steinbach, P. J., Eaton, W. A., Szabo, A. & Hofrichter, J. Effects of chain stiffness on the dynamics of loop formation in polypeptides. Appendix: testing a 1-dimensional diffusion model for peptide dynamics. The Journal of Physical Chemistry B 106, 11628–11640 (2002).

6. Faesen, A. C. et al. Basis of catalytic assembly of the mitotic checkpoint complex. Nature 542, 498–502 (2017).

7. Piano, V. et al. CDC20 assists its catalytic incorporation in the mitotic checkpoint complex. Science 371, 67–71 (2021).

8. Roy, B., Verma, V., Sim, J., Fontan, A. & Joglekar, A. P. Delineating the contribution of Spc105-bound PP1 to spindle checkpoint silencing and kinetochore microtubule attachment regulation. Journal of Cell Biology 218, 3926–3942 (2019).

9. Banerjee, A., Chen, C., Humphrey, L., Tyson, J. J. & Joglekar, A. P. BubR1 recruitment to the kinetochore via Bub1 enhances Spindle Assembly Checkpoint signaling. Molecular Biology of the Cell, In press.

10. Alfonso-P’ serez, T., Hayward, D., Holder, J., Gruneberg, U. & Barr, F. A. MAD1-dependent recruitment of CDK1-CCNB1 to kinetochores promotes spindle checkpoint signaling. Journal of Cell Biology 218, 1108–1117 (2019).

11. Li, J., Gao, J.-Z., Du, J.-L., Huang, Z.-X. & Wei, L.-X. Increased CDC20 expression is associated with development and progression of hepatocellular carcinoma. International Journal of Oncology 45, 1547–1555 (2014).

12. Nilsson, J., Yekezare, M., Minshull, J. & Pines, J. The APC/C maintains the spindle assembly checkpoint by targeting Cdc20 for destruction. Nature Cell Biology 10, 1411–1420 (2008).

13. Kafle, B. P. Chemical Analysis and Material Characterization by Spectrophotometry (Elsevier, 2020).

14. Heinrich, S. et al. Determinants of robustness in spindle assembly checkpoint signalling. Nature Cell Biology 15, 1328–1339 (2013).

